# Antimicrobial Susceptibility Profiles of Bacteria Isolated from The Ozama River in Santo Domingo, Dominican Republic

**DOI:** 10.1101/2022.09.19.508624

**Authors:** Roberto Bonnelly, Ana Lídia Q. Cavalcante, Camila Del Rosario Zorrilla, Victor V. Calderón, Albert Duarte, Rafael Baraúna, Rommel T. Ramos, Yaset Rodriguez, Luis E. Rodriguez de Francisco, Luis O. Maroto, Omar P. Perdomo, Edian F. Franco

## Abstract

The dissemination of antimicrobial-resistant bacteria in environmental waters is an emerging concern in medical and industrial settings. In the present study, our research team analyzed superficial water samples from 3 different collection sites along the Ozama River, the most important river in the Dominican metropolitan area. Seventy-six isolates were obtained from culture media previously enriched with cefotaxime and imipenem and subsequently identified by MALDI-ToF. Our isolates spanned 12 genera of bacteria; over 30% were of clinical relevance, and 43% exhibited a phenotype classified as multi-drug resistance. The most frequent species identified as *Stenotrophomonas maltophilia* (n = 33), an emerging nosocomial pathogen. This study constitutes part of the initiative to understand the profiles of the perils of multi-drug resistance in metropolitan areas of the Dominican Republic: a nation with poor antibiotic use regulation.

**IMPORTANCE:** Nearly two billion people are sourcing their water from low-quality reservoirs world-wide. These reservoirs consist of contaminated waters with anthropogenic pollutants such as antibiotics, disinfectants, and other substances used to treat water in societies with scarce resources and unregulated industries. Furthermore, the exposure of these antibiotics to potable water reservoirs promotes the increase of clinically relevant bacteria with antibiotic-resistance capabilities, becoming a public health crisis. Therefore, treating patients with infectious diseases and providing prophylactic measures against infection-associated conditions (such as post-surgical recovery) has become progressively more difficult. Based on this evidence frame, it is of considerable importance to analyze the bacterial profiles of rivers that fall prey to anthropogenic contamination, as these investigations on antibiotic resistance will, of course, benefit the life of humans.

## INTRODUCTION

New antibiotics have been developed over the last fifty years to ease the increasing need for effective antimicrobial prophylaxis and therapies. The adoption of antibiotics to prevent and treat infections in the animal husbandry industry and their use at sub-therapeutic concentrations are growth promoters for agriculture (1, 2, 3, 4, 5). As a result, they have propelled their abundance in the environment (6). However, if performed irresponsibly, these practices lead to inadequate disposal of antibiotics and other pollutants directly to essential water sources. These deposits, especially in aquifer systems of developing countries, contribute to the rise of drug-resistant bacteria in environmental reservoirs (7, 8, 9), increasing antibiotic resistance (AR) in clinically relevant bacteria (10). For this reason, the emergence and spread of antibiotic-resistant bacteria (ARB) and antibiotic-resistant genes (ARG) has become a significant public health crisis worldwide (11).

Due to the increase in antibiotic prescription and misuse, aquatic systems are now considered critical reservoirs of these genetic elements (2). However, this is not only due to the misuse of antibiotics; recent studies have demonstrated that these same aquatic environments accumulate substantial amounts of contaminants (such as fecal matter, metals, disinfectants, and hormones) (12). These same contaminants are involved in selecting resistant bacteria in the environment (13). As a result, antibiotic-resistant bacteria reside in multiple aquatic environments (14). Examples of these are *Klebsiella pneumonia, Enterococci spp, Pseudomonas spp* which is consistently found as a multi-drug resistant bacteria across many studies and is considered to be extremely detrimental to patients (5). Multi-drug resistant colonies of these bacteria are now commonly found in environmental waters, now as emergent contaminants, themselves (15).

Consequently, ARBs are one of the greatest threats to public health in the 21st century, as described by the World Health Organization (16). Therefore, international large-scale and local monitoring systems are urgently needed to assess their occurrence in the environment, primarily in water bodies that are essential sources for the survival of humanity. This study unveils the multi-drug resistance prevalence and microbiome characteristics of the Ozama River in the Dominican Republic.

The Ozama River is one of the main drainage basins of the Dominican Republic, covering an approximate area of 2,847.15 km2, traversing rural and urban areas and finally discharging its waters into the Caribbean Sea. The river’s area of influence has a population density of approximately 3.8 million people (17, 18, 19). Since colonial times (late 15th century for La Hispaniola), the Ozama River has been a symbol of prosperity because of its magnitude, becoming the most important river in the Dominican Republic. On its margins, the city of Santo Domingo was born, the first city in America founded in 1498, and housed many critical historical artifacts (20). The river as a resource was pivotal for the socio-economic development of Santo Domingo; its waters serve for agriculture, fishing, recreation, and potabilization for the metropolitan area. It is also crucial for industrial activity and residential discharge, such as metal industries, pharmaceutical factories, and residential houses (20, 21).

The Ozama River has experienced a growth in demand due to Santo Domingo’s demographic increase. This increase has caused the occupation of its river banks with inadequately prepared settlements and industries, which in turn generates high levels of physical, chemical, and biological contamination to the River (20, 21, 19). The river receives discharges of at least 54 sewage waterways, which transport sanitary wastes to the river, both directly and indirectly. In addition to its banks, approximately 241 companies and industries dump their liquid waste into the river (22, 23). Also, the river receives large amounts of solid waste (garbage) discarded by the settlements on its banks; according to (21), the Ozama river receives, as many other urban rivers, receives important amounts of garbage every year (24).

The river suffers from abnormally high anthropogenic environmental impact, causing the proliferation of water hyacinth, an invasive aquatic plant that develops in highly contaminated water basins (25). Most of the contamination comes from organic matter, which causes the excessive proliferation of bacteria in these waters. The river has been studied directly in the basin or its secondary tributaries, where the presence of pathogenic bacteria is notorious. One study by (26) found high levels of coliforms present in the waters, where they managed to isolate *Escherichia coli, Klebsiella spp., Pseudomonas aeruginosa, Enterobacter cloacae, Proteus spp*. and *Salmonella spp*. among others. These bacteria presented their highest prevalence at the confluence of the Ozama and Isabela rivers.

The Ozama river has many tributaries; the Cabon River, researched by (27), where the microbiological tests found concentrations of fecal coliforms above the maximum allowed by the Dominican Standard Surface Waters Norm. The Ozama river’s coliform count was estimated to be 70 - 70,000 CFU/100 ml in 44% of the samples obtained from the river; in the other 56%, the values were untouchable due to high microbial prevalence. This study also detected *Streptococcus spp., Salmonella spp, Shigella spp, Klebsiella spp, Alteromonas spp, Enterobacter spp*, and *Pseudomonas spp* in the Ozama river samples.

A previous study (6), was examined the dissemination of resistance to beta-lactams in the bacterial communities present in the Isabela River, the main tributary of the Ozama River. This tributary is also highly impacted by anthropogenic action. This study found that the most common genera in the Ozama’s main tributary were *Acinetobacter spp*. (44.6%) and *Escherichia spp*. (18%). We also identified twenty clinically-relevant bacteria isolated in urban areas of the tributary; these isolates presented the following genes: KPC-3, OXA-1, OXA-72, OXA-132, CTX-M-55, CTX-M-15, and TEM-1. All of these genes are directly responsible for antibiotic resistance to beta-lactams.

This work aims to describe the antimicrobial susceptibility profiles of the bacterial communities present at three points with different anthropogenic impacts in the Ozama River. This study will deliver results through microbiology techniques and the sequencing of the genomes of the isolates that present higher levels of resistance.

## RESULTS

### Physical, Chemical, and Biological Parameters of Water

We performed a series of analyses described by the Environmental Standard for Surface and Coastal Zones Water Quality (28) for acceptable physical, chemical, and biological limits. For this study, we classified surface waters based on the current or potential use of its waters: Class A, these waters are suitable for vegetable irrigation, recreational uses with direct contact (like swimming), and human and animal consumption without previous treatment; and class B waters which could serve as a public drinking water supply, irrigation of crops, industrial uses, and livestock maintenance given adequate treatment(28). Our research team sampled the Ozama River in June 2019; this pre-dates the current pandemic, which will give us a baseline for our subsequent surveillance studies. Due to our sampling nature, we did not assess seasonal microbiome profile variation. However, it is fundamental to note the consistent temperature in these regions, as they usually fluctuate between 30°C and 35°C throughout the year. During our sampling, we did not record any rains.

Our sampling strategy and naming scheme consisted of three primary regions; we named them: point A, corresponding to Cañada la Rubia, an area where a highly populated zone discharges waste; B, the river delta, near the confluence with Isabela River (6); and C, near its source. Each point was sampled three times, corresponding to replicas one, two, and three.

Our team moved to these in-river locations via a small motorboat. Once well positioned, we proceeded to take samples: Primarily, we conditioned each recipient with river water. Next, we used a weighted sampling bottle to collect water three meters below the surface. This strategy was repeated twice around the same area, no more than 30 meters away from the first sampling point.

Our assays consisted of nine measurements, one microbiological and eight physicochemical, based on the Environmental Standard for Surface and Coastal Zones Water Quality. We described these assays in Table 1 (Supplementary data 1: the complete list of measured parameters).

**TABLE 1.**
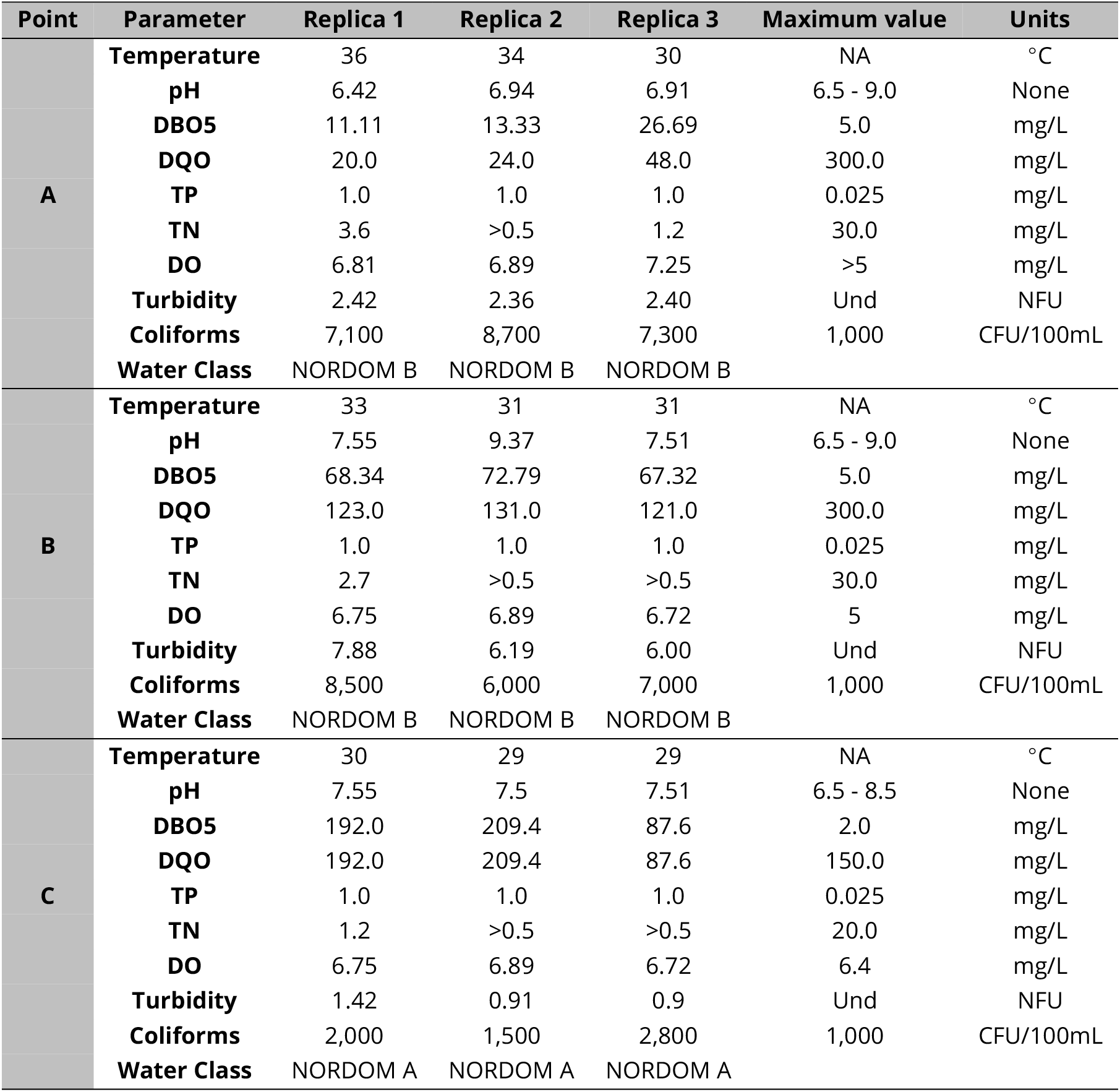
Physicochemical and microbiological results of water sampled from Ozama River.

| Point | Parameter | Replica 1 | Replica 2 | Replica 3 | Maximum value | Units |
| --- | --- | --- | --- | --- | --- | --- |
| **A** | **Temperature** | 36 | 34 | 30 | NA | °C |
|  | **pH** | 6.42 | 6.94 | 6.91 | 6.5 - 9.0 | None |
|  | **DBO5** | 11.11 | 13.33 | 26.69 | 5.0 | mg/L |
|  | **DQO** | 20.0 | 24.0 | 48.0 | 300.0 | mg/L |
|  | **TP** | 1.0 | 1.0 | 1.0 | 0.025 | mg/L |
|  | **TN** | 3.6 | >0.5 | 1.2 | 30.0 | mg/L |
|  | **DO** | 6.81 | 6.89 | 7.25 | >5 | mg/L |
|  | **Turbidity** | 2.42 | 2.36 | 2.40 | Und | NFU |
|  | **Coliforms** | 7,100 | 8,700 | 7,300 | 1,000 | CFU/100mL |
|  | **Water Class** | NORDOM B | NORDOM B | NORDOM B |  |  |
| **B** | **Temperature** | 33 | 31 | 31 | NA | °C |
|  | **pH** | 7.55 | 9.37 | 7.51 | 6.5 - 9.0 | None |
|  | **DBO5** | 68.34 | 72.79 | 67.32 | 5.0 | mg/L |
|  | **DQO** | 123.0 | 131.0 | 121.0 | 300.0 | mg/L |
|  | **TP** | 1.0 | 1.0 | 1.0 | 0.025 | mg/L |
|  | **TN** | 2.7 | >0.5 | >0.5 | 30.0 | mg/L |
|  | **DO** | 6.75 | 6.89 | 6.72 | 5 | mg/L |
|  | **Turbidity** | 7.88 | 6.19 | 6.00 | Und | NFU |
|  | **Coliforms** | 8,500 | 6,000 | 7,000 | 1,000 | CFU/100mL |
|  | **Water Class** | NORDOM B | NORDOM B | NORDOM B |  |  |
| **C** | **Temperature** | 30 | 29 | 29 | NA | °C |
|  | **pH** | 7.55 | 7.5 | 7.51 | 6.5 - 8.5 | None |
|  | **DBO5** | 192.0 | 209.4 | 87.6 | 2.0 | mg/L |
|  | **DQO** | 192.0 | 209.4 | 87.6 | 150.0 | mg/L |
|  | **TP** | 1.0 | 1.0 | 1.0 | 0.025 | mg/L |
|  | **TN** | 1.2 | >0.5 | >0.5 | 20.0 | mg/L |
|  | **DO** | 6.75 | 6.89 | 6.72 | 6.4 | mg/L |
|  | **Turbidity** | 1.42 | 0.91 | 0.9 | Und | NFU |
|  | **Coliforms** | 2,000 | 1,500 | 2,800 | 1,000 | CFU/100mL |
|  | **Water Class** | NORDOM A | NORDOM A | NORDOM A |  |  |

Based on our results, we concluded that sampling points A and B can be classified as Class B Water Quality. We also provided evidence suggesting point C corresponds to a Class A Water Quality. However, we also encountered evidence for all three points suggesting contamination: our assays revealed that Point C had considerably altered (25% higher) chemical oxygen demand (COD) than recommended standards and higher than our two other sampling sites. We also evidence higher biological oxygen demands (BOD) for these points, as they were over two orders of magnitude above recommended values for one sample. Our total coliform concentration also at least twice the recommended values for most samples.

### Composition and Distribution of the Isolates

Our isolation methodology consisted of an initial antibiotic-based selection and culturing in differentiation media. First, we performed the selection phase with MacConkey agar supplemented with antibiotics: cefotaxime (CTX), and imipenem (IMP), to screen for resistance. This methodology yielded seventy-six isolates. Next, our team performed colony isolation using chromogenic agar. Through this method, we recovered forty isolates from media initially selecting CTX resistance and thirty-six isolates from IMP media. Finally, we isolated most of our specimens from Point B, totaling twenty-eight colonies, while Point A and Point C combined yielded twenty-four isolates.

The most frequent genus in isolates was *Stenotrophomonas spp* (46%), the predominant species was *Stenotrophomonas maltophilia* (n=33), followed by *Pseudomonas spp* (23%), with *Pseudomonas protegens* being the most abundant species (n = 4), the Acinetobacter genus (22%) with the highest presence of *Acinetobacter baumannii* (n = 7), then the *Klebsiella* genus (5%), with the *Klebsiella pneumoniae* (n = 4), and Escherichia genus (4%) with *Escherichia coli* (n = 3), being the least number of isolates found (Figure 1).

**FIG 1.**
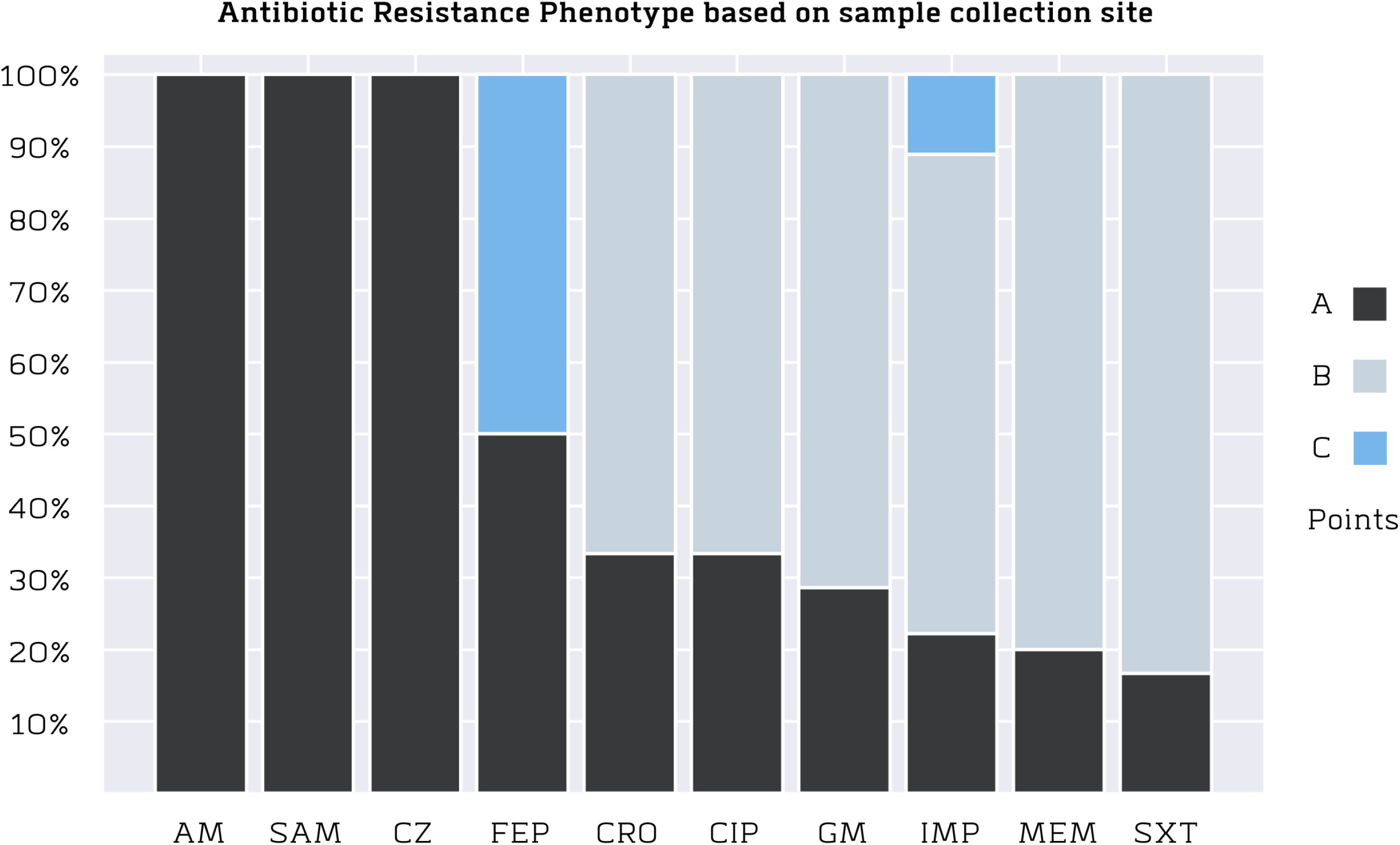
Distribution of the most frequent bacterial genera found in all points of the Ozama River.

The region that presented the most variety was the one corresponding to the Rio Delta (Point 2) with five genera, in which the most abundant were the genera *Stenotrophomonas* (47%) and *Pseudomonas* (21%), standing out for the presence of the genus *Klebsiella* (14%). The Cañada de la Rubia region (Point 1) presented 4 genera, of which the most abundant genera were *Stenotrophomonas* (50%) and *Pseudomonas* (25%). The Nacimiento del río (Point 3) presented only three genera corresponding to *Stenotrophomonas* (42%), *Pseudomonas* (25%), and *Acinetobacter* (33%), this last genus was the most abundant at this point compared to its presence in the Points 1 and 2 (Figure 2).

**FIG 2.**
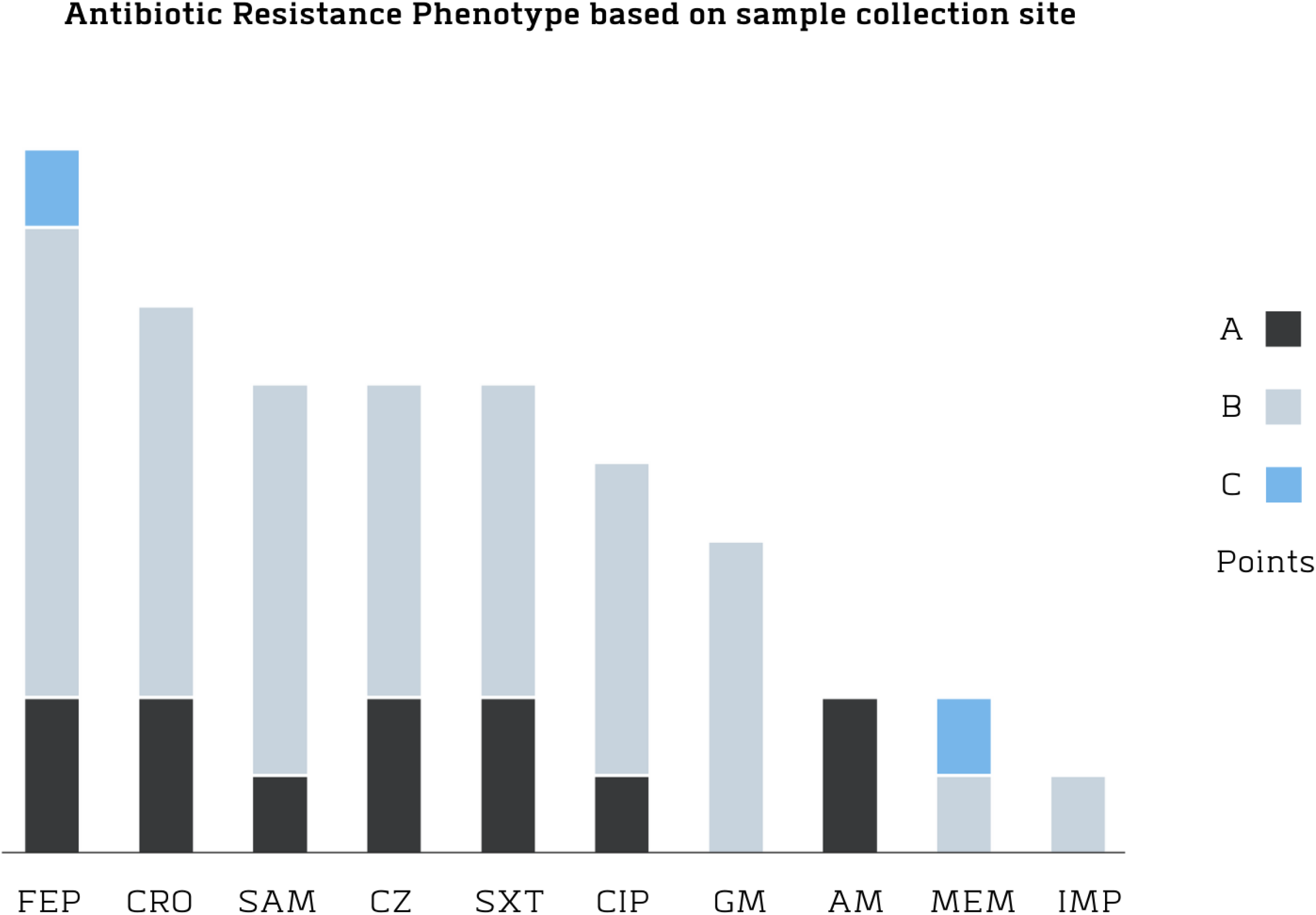
Distribution of the bacteria genera present in each of the monitored points in the Ozama river basin.

### Antibiotic Resistance

Twenty-three isolates, a total of 10 (44%), presented resistance. However, only seven presented resistance to 3 or more of the 14 groups of antibiotics, considering themselves to be a multi-resistant phenotype (Table 2). Klebsiella (five isolates) and Escherichia (one isolate) presented substantial phenotypic variety; we measured resistance to seven antibiotics, especially cephalosporins. Also, these isolates tested positive for broad-spectrum beta-lactamase genetic markers. We found similar resistance diversity to those presented in previous studies based on antibiotic resistance assays performed with ciprofloxacin, gentamicin, and trimethoprim/sulfamethoxazole (6).

**TABLE 2.** Resistance phenotype of the different isolates of clinical interest obtained from the Ozama river

| ID | Species | Resistance phenotype |
| --- | --- | --- |
| DC1 | *Acinetobacter baumannii* | No resistance |
| DC2 | *Escherichia coli* | AMP, CZ, FEP, CRO, SXT |
| DC5 | *Acinetobacter baumannii* | No resistance |
| DC8 | *Escherichia coli* | AMP, SAM, CZ, FEP, CRO, CIP, SXT |
| DC9 | *Acinetobacter baumannii* | No resistance |
| DC10 | *Acinetobacter pittii* | No resistance |
| EC1 | *Klebsiella pneumoniae* | SAM, CZ, FEP, CRO, CIP, GM, SXT |
| EC4 | *Klebsiella pneumoniae* | SAM, CZ, FEP, CRO, CIP, GM, SXT |
| EC6 | *Escherichia coli* | SAM, CZ, FEP, CRO, CIP, GM, SXT |
| EC7 | *Klebsiella pneumoniae* | SAM, CZ, FEP, CRO, CIP, GM, SXT |
| EC11 | *Acinetobacter baumannii* | No resistance |
| EC13 | *Acinetobacter baumannii* | No resistance |
| EC14 | *Acinetobacter baumannii* | FEP, IPM |
| EC15 | *Acinetobacter baumannii* | MEM |
| EC16 | *Acinetobacter baumannii* | No resistance |
| EC16.2 | *Klebsiella pneumoniae* | SAM, FEP, CRO |
| FC1 | *Acinetobacter pittii* | No resistance |
| FC4 | *Acinetobacter baumannii* | No resistance |
| FC5 | *Acinetobacter baumannii* | No resistance |
| FC6 | *Acinetobacter baumannii* | No resistance |
| FC7 | *Acinetobacter baumannii* | FEP, MEM |
| FC8 | *Acinetobacter baumannii* | No resistance |
| FC12 | *Acinetobacter baumannii* | No resistance |

Abbreviation: Amikacin, AN; Ampicillin, AMP; Ampicillin/Sulbactam, SAM; Cefazolin, CZ; Cefepime, FEP; Ceftazidime, CAZ; Ceftriaxone, CRO; Ciprofloxacin, CIP; Ertapenem, ETP; Gentamicin, GM; Imipenem, IPM; Meropenem, MEM; Piperacetyline, TZP; Trimethoprim-Sulfamethoxazole, SXT.

The phenotypes described in our study indicate that the highest level of resistance to multiple antibiotics is at Point 2, corresponding to the region of the River Delta. The abundance of resistance to ampicillin/sulbactam, cefazolin, ceftriaxone, gentamicin, and trimethoprim/sulfamethoxazole was more than two times greater than in Point 1; therefore, there exists the possibility of positive selection for these phenotypes. Our results show no significant resistance patterns to most antibiotics, except for cefepime and meropenem (Figure 3).

**FIG 3.**
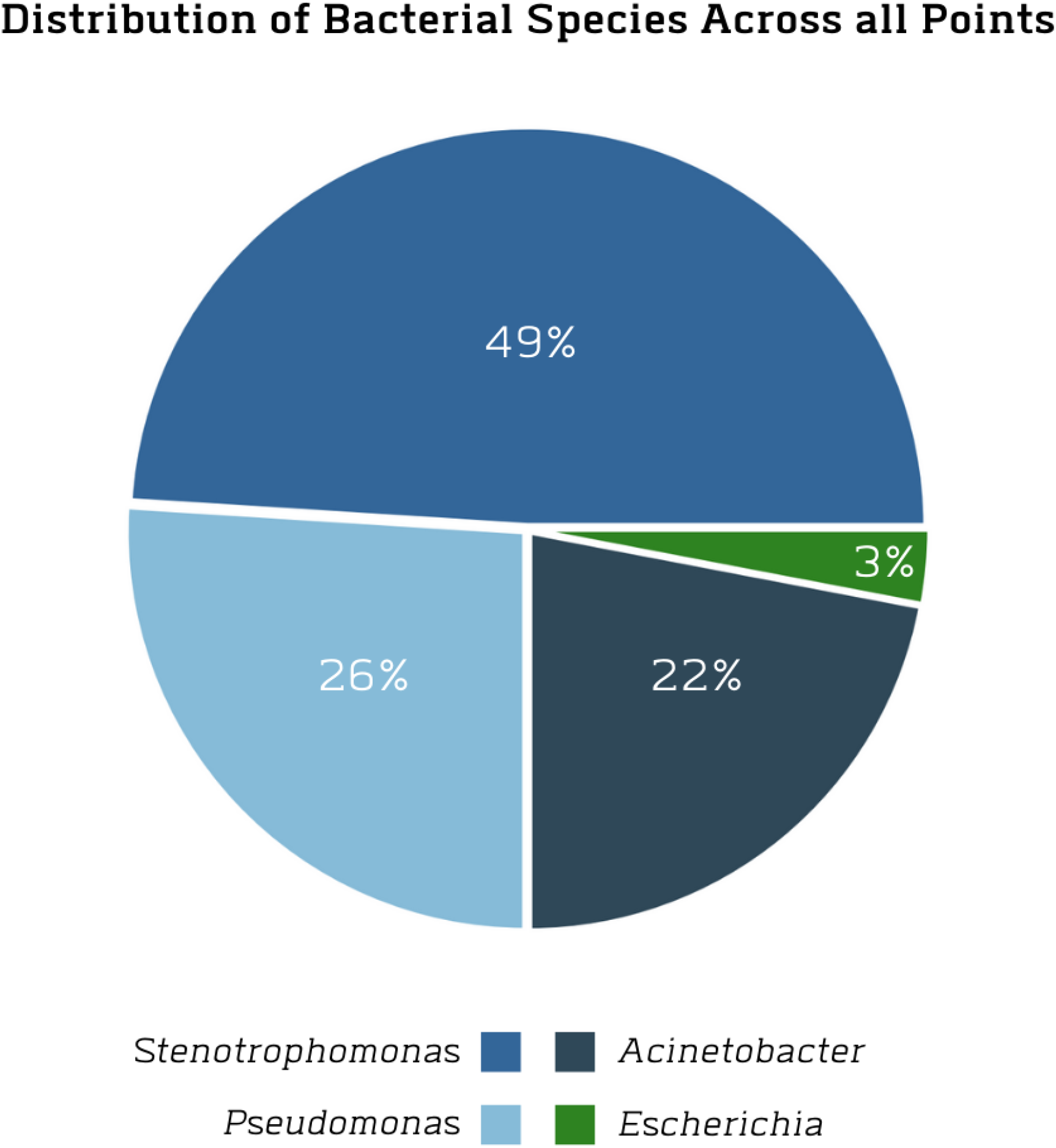
Distribution of resistant isolates at different sampling points.

### Analysis of Multi-Resistance Genomes

In order to describe resistomes found in these waters, we proceeded with whole genome sequencing of our isolates. We sequenced and assembled seven genomes. Our pipeline consisted of the following steps: quality control, data pruning, and de Bruijn graph assembly. Our team built an in-house assembly pipeline for this analysis. We named our genomes DC2, DC8, DC10, EC4, EC7, FC5, and FC7, as displayed in our table 3.

**TABLE 3.** Major genomic characteristics of genomes obtained from Ozama river.

| Sample ID | Species | Genome Size (pb) | GC (%) | CDS | N50 | Plasmids |
| --- | --- | --- | --- | --- | --- | --- |
| DC8 | Escherichia coli | 4,710,092 | 50.7 | 4,664 | 148,992 | IncY, IncFIB |
| DC2 | Escherichia coli | 4,657,304 | 51.1 | 4,637 | 56,323 | IncY |
| DC10 | Acinetobacter pittii | 3,907,124 | 38.6 | 3,705 | 245,032 | - |
| EC4 | Klebsiella pneumoniae | 5,527,705 | 57.2 | 5,457 | 201,357 | IncFBI(K), IncFII(K), Col440II |
| EC7 | Klebsiella pneumoniae | 5,424,764 | 57.3 | 5,346 | 320,811 | IncFBI(K), IncFII(K0), Col440II |
| FC5 | Acinetobacter baumannii | 3,809,530 | 38.9 | 3,622 | 324,095 | - |
| FC7 | Acinetobacter baumannii | 3,841,875 | 38.9 | 3,660 | 323,883 | - |

#### Escherichia coli

Two *E. coli* isolates were sequenced (DC2 and DC8). Some of the computed parameters were the genome size, GC% content, and total predicted coding sequences (CDS). The isolates presented an average genome size of 4.6 Mb with a GC% content of 51%, and between 4,637 and 4,664 CDS as seen in table 3.

A total of 107 CDSs were related to virulence, disease, and defense. PlasmidFinder1.1 found IncY plasmids sequence, and VirulenceFinder2.0 results presented a significant amount of virulence factors, including terC, traT, neuC, capU, ipfA, and iss. Among the Antibiotic Resistance Genes (ARGs) found in the sequences with perfect hits were msbA, acrB, acrA, mdtH, QnrS1, marA, tolC, mdtG, dfrA12, aadA2, sul1, baerR, and H-NS. They presented resistance to antibiotic classes like penem, sulfonamide, fluoroquinolone, and carbapenem. The ARG mechanisms found were antibiotic efflux, antibiotic target alteration, antibiotic inactivation, antibiotic target protection, antibiotic target replacement, and reduced antibiotic permeability, as seen in table 2. Serotype classification for these genomes was O104, O8, O9, O9a, H11, and H30. According to PathogenFinder1.1, these isolates have a probability of 94% of being human pathogens.

#### Pseudomonas putida

One Pseudomonas was sequenced. The isolate presented a genome size of 5.5 Mb with a GC% content of 63.4%, and a total of 5234 CDS as seen in table 3. Approximately 1% of the genes were related to virulence, disease, and defense. In addition, there were found resistances to fluoroquinolone and tetracycline in the resistome. Some of the identified genes were adeF and Acinetobacter baumannii abaQ. PathogenFinder1.1 calculated that this isolate has a 17.4% probability of being a human pathogen. This genome did not present any mobile genetic element (MGE).

#### Acinetobacter pittii

Only one isolate was obtained. This isolate contained a 3.9 Mb size with a 38.6% of GC content and 3,705 CDS, where 43 were related to virulence, disease, and defense. According to the Resistance Genes Identifier, only nine strict hits were identified, these were ADC-15, OXA-421, adeF, abaQ, parC, AmvA, AbaF, and LpsB. Resistance to fluoroquinolone, tetracyclin, cephalosporin, macrolide, carbapenem, penam, aminocoumarin, Fosfomycin, and acridine dye was found. The resistance mechanism detected included antibiotic efflux, antibiotic inactivation, reduced permeability to antibiotics, and antibiotic target alteration. According to PathogenFinder1.1, this isolate had a 92.5% probability of being a human pathogen. No MGE were found in this genome.

#### Klebsiella pneumoniae

We sequenced two *K. pneumoniae* isolates. These genomes had an average size of 5.47 Mb with an average of 57.25% of GC content. Also, the CDS went between 5346 and 5,457, where 136 were related to virulence, disease, and defense. There were nine perfect hits and 30 strict hits in the resistome identifier. Among the perfect hits, there were resistances to fluoroquinolone, glycylcycline, tetra-cyclin, carbapenem, diaminopyrimidine, nitrofuran, aminocoumarin, cephalosporin, penam, sulfonamide, penem, monobactam, rifamycin, and macrolide. Among the identified genes, oqxA, LptD, KpnF, SHV-28, sul2, TEM-1, CTX-M-15, QnrB1, and OXA-1. The resistance mechanism included antibiotic efflux, antibiotic inactivation, antibi-otic target replacement, antibiotic target protection, antibiotic target alteration, and reduced antibiotic permeability. Among the mobile genetic elements, there were Inc and Col plasmids sequences. IncFIB(K), IncFII(K), and Col440II were found. According to PathogenFinder2.0, these genomes had approximately 89.4% probability of being a human pathogen. Finally, there were virulence factors identified by VirulenceFinder2.0 that included iutA, traT, fyuA and irp2.

#### Acinetobacter baumanii

Two A.baumanii isolates were obtained. These genomes had an average size of 3.8 Mb with an average GC content of 65%. The CDS were between 3660 and 5346, where approximately 115 were related to virulence, disease, and defense. According to the Resistance Genes Identifier, there were four perfect hits and 12 strict hits. Among the perfect hits there were adeL, abeS, adeK, and adel. These genes included resistances to fluoroquinolone, tetracycline, macrolide, aminocoumarin, lincosamide, carbapenem, cephalosporin, rifamycin, diaminopyrimidine, phenicol, and penem. The resistance mechanisms of these genomes included antibiotic efflux, antibiotic inactivation, antibiotic target alteration, and reduced permeability to the antibiotic. According to PathogenFinder1.1, these isolates had approximately 88.5% probability of being a human pathogen. These isolates had no mobile genetic elements identified by PlasmidFinder2.0.

## DISCUSSION

The prolonged interactions between the Ozama River, its communities, and surrounding industries evolved into a harmful reservoir for antibiotic resistance in the Dominican Republic. Although this interaction was crucial to the development of the city of Santo Domingo, prolonged anthropogenic influences from the surrounding population are now evident through contamination. During colonial times, its waters served as a transportation mechanism and a source of potable water and fish for consumption. Also, this river was essential as a wastewater dump for its residents. However, the ever-increasing population surrounding the perimeter of the river has become a public health and sanitary challenge, as delivering treated water properly to these settlements has not been possible.

We must also mention the ongoing industrialization of the river’s banks, including metal refineries, abattoirs, hospitals, and residential zones with significant pollution production. As many researchers have pointed out in previous publications, the continued integration of untreated wastewater into an ecosystem is a primary motivation for altering antimicrobial resistance profiles. The disruption of these ecosystems with the introduction of antimicrobial agents most like will result in the expansion of resistance genes.

Perturbation in these environments can result in a higher prevalence of antibiotic-resistant bacteria and resistance genes; these modifications may pose a significant danger to human and environmental health. Our study explored and described the antimicrobial resistance composition in three contrasting areas of the Ozama River. Our primary focus was beta-lactamase-producing gram-negative bacteria of clinical relevance.

### Physico-chemical and Microbiological Analyses

Our research team performed physicochemical analyses as the Standard Methods for the Examination of Waters and Wastewater dictated. Our results indicate a critical rise in average biological oxygen demand (BOD) across all sampling points. Furthermore, these levels of BOD are correlated to the large populations surrounding the river, as these settlements do not have access to a proper sewage system, hence dumping all of their wastewater into the river. It is also essential to mention the presence of active abattoirs without a proper wastewater treatment process on their banks.

Many restoration projects have taken a course on the Ozama River. Among them is OceanCleanup’s Interceptor 004 initiative, a very ambitious project to remove debris from the river. However, a proper follow-up to this project could be implementing more strict policies around antibiotics and their use. Nowadays, all families can buy antibiotics directly from any pharmacy without a prescription. The frictionless acquisition of antibiotics by uneducated patients exacerbated our nation’s position against antibiotic resistance during the pandemic, as these individuals attended to their viral infections and preventive needs with antibiotics. **The deleterious impact of this popular practice has been stressed by many research in the last years** [citas X1, X2]

Our isolates and their peculiar distribution are not that surprising, as these are the same bacteria commonly found in infected patients at hospitals around the city, especially in low-income districts and neighborhoods. Infections by *Acinetobacter, Escherichia*, and *Pseudomonas*, are among the most common in clinical settings, and their resistance phenotypes can only suggest a continuous interaction with antibiotics.

Of all the isolates, those particularly dangerous in hospitals, nursing homes, and patients whose care requires devices such as ventilators and intravenous catheters (29) (*Klebsiella, Escherichia*, and *Acinetobacter*) have undergone sensitivity tests. The genus *Stenotrophomonas* was not considered for testing because it is intrinsically multi-resistant despite being an opportunistic human pathogen (30). Considering the distribution of our isolates, we compared their distribution to the National Laboratory of Public Health’s, we found that both *Klebsiella pneumoniae* and *Escherichia coli* are presented as the most common pathogens.

### Genomic and Resistome Analyses

Our genomic analyses allowed us to identify our isolates through phylogenetics and discover the virulence factors, antibiotic-resistance genes, and essential features they harbor. These analyses mainly consisted of reconstructing each individual’s genome to extract coding regions and compare them to known antibiotic resistance genes through genome alignment and annotation.

Our pipeline effectively identified regions of interest in our samples, ranging from ARG-harboring plasmids to pathogenic toxins and other antibiotic-resistance genomic features. Our main finding through this methodology is the extraordinary amount of pathogenic bacteria harboring antibiotic-resistance genes.

Isolate DC8, an *Escherichia coli* found in waters close to many residential zones harbors resistance genes against beta-lactamases, fluoroquinolone, and cephalosporins. As do isolates EC1, EC4, EC6, and EC7, all recovered from the Ozama river’s delta.

Isolate EC14, an *Acinetobacter baumannii* from the river’s delta, presented resistance to beta-lactamases, specifically carbapenems, a last-resort antibiotic. It is important to note that *A. baumannii* is an opportunistic pathogen, mainly causing nosocomial infections.

We also detected plasmids as part of our pipeline: DC8 harbored plasmids IncY and IncFIB. IncY commonly carries the antibiotic resistance gene CTX-M-15, a serine beta-lactamase, more specifically, a cefotaximase. This ARG is one of the most common causes of resistant infections worldwide. The other plasmid, IncFIB, is commonly associated with gene blaIMP, which transfers resistance to last-resort carbapenem antibiotics.

As we described, it is crucial to properly attend to these genes, plasmids, and other genomic features in the urban waters of Santo Domingo. Furthermore, inadequate wastewater disposal into this river can and ultimately will cause severe deterioration to the population and settlements that depend on this water resource. Therefore, it is paramount that we, the researcher, notify appropriate authorities of our results and be of assistance in generating new policies regarding monitoring, assessment, and continuous care of our environment and water supply.

## MATERIALS AND METHODS

### Sampling and Physico-chemical tests

Ourteam sampled the rivers in June 2019, prior to the SARS-CoV-2 pandemic. Our main goal was to sample the Ozama River (18°28’0.12”N, 69°52’59.88”W) in three distinct states: pristine (untouched by humans), moderate human intervention (near a settlement), and considerably intervened (near wastewater disposal conduits). For the first state, we sampled the river source in “Loma Siete Cabezas, Sierra de Yamasá” which covers 148 km and empties into the Caribbean Sea. We collected one liter of water for each of the three samples collected from this point and made triplicates for microbiological and physicochemical analyses. We repeated this process for every point we sampled. We chose a community upstream of the city as the moderately intervened point and a wastewater drainage output for the most contaminated. The bottles were kept in an icebox and transported to the microbiology and Physical-chemistry laboratories within 6h of collection.

We measured pH, Chemical Oxygen Demand (COD), Biological Oxygen Demand (BOD), Dissolved oxygen, turbidity, total phosphorous, and total nitrogen were the chemical-physical parameters determined, following the procedures described in the book “Standard Methods for the Examination of Water and Wastewater”. We also measured total coliforms in all of our samples.

### Bacteria Isolation

Aliquots of 1, 10, and 50 mL were filtered from the samples using the standardized membrane filtration method using 0.22μm diameter cellulose filter (Millipore). The membranes were placed on 2 MacConkey agar media (Oxoid, UK), one with 4 g/mL of imipenem and the other with 8 g/mL of cefotaxime, incubated at 37°C for 24-48 hours, following the recommendation of (7). The individual colonies were purified on the same media and stored in 25% glycerol at −70°C. Then, the individual colonies from this process were isolated in chromogenic culture media (ChromoAgar) and stored in 25% glycerol at −70°C.

### Bruker BioTyper Bacterial Identification

About 0.1 mg of each culture was inoculated into a kit sample carrier (MPS 96 target polished steel), then the samples were coated with 1L of matrix solution consisting of cyano-4-hydroxycinnamic acid in 50% acetonitrile and 2.5% trichloroacetic acid and allowed to dry at 25°C for 15 minutes. The identification of the isolated strains was performed with the MALDI-TOF (matrix-assisted laser desorption ionization-time of flight) technique (29), with the BioTyperR©3.1 software (Bruker Daltonics, Germany) equipped with the MBT 6903 MPS library (2019), using the MALDI BioTyper Standard Processing Method and the MALDI Biotyper MSP Standard Identification Method adjusted by the manufacturer.

### Antibiotic Susceptibility Test

The selected strains were replicated in chromogenic culture (ChromoAgar) and incubated at 37°C for 24 hours. The susceptibility test was performed based on the minimum inhibitory concentration (MIC) and then classified according to the recommendations of the Institute of Clinical and Laboratory Standards (30).

BD Phoenix ID broth and AST broth were used for sample preparation, in conjunction with the BD PhoenixTM AP kit, where the inoculum was standardized from 0.25 to 0.5 according to the required McFarland. The broths with the samples were placed on a BD PhoenixTM NMIC-406 panel for gram negatives, and the automated identification and susceptibility test systems BD PhoenixTM 100 and BD PhoenixTM M50 with the data management system BD EpiCenterTM, were the susceptibility test was carried out at a time interval of 12 to 13 hours. Isolates were treated with the following antibiotics: ampicillin, amikacin, amoxicillin-clavulanic acid, ceftazidime, cefuroxime, ciprofloxacin, gentamicin, imipenem, trimethoprim/sulfamethoxazole, meropenem, and ertapenem.

### Data Analysis

The physicochemical parameters were compared with the Quality Standard for Surface Waters and Coastal Zones established by the Ministry of Environment and Natural Resources to determine possible relationships between the levels of contamination and the bacteria found. The data was processed through component analysis to identify similarities between sampling sites.

### Genomic DNA Extraction From Isolates

Genomic DNA was extracted from colonies incubated in TSB for 24 h at 35° C. One aliquot of 4 mL of culture was centrifuged at 8,000 g for 2 min. Cell pellet was subjected to the QIAamp DNA extraction kit (Qiagen, Germany) with the adaptations indicated next: the bacterial pellet was suspended in 420 μL of the modified lysis buffer (20 *μL* proteinase K, 200 *μL* of TSB, and 100 *μL* of Qiagen’s ATL buffer), and incubated for 10 min at 56° C. The addition of 50 μL of absolute ethanol followed by 3 min incubation at room temperature concluded the adaptations; from this point in the process, the protocol continued according to the manufacturer’s recommendations. DNA obtained was suspended in 50 μL of Qiagen’s TE buffer. The integrity of extracted DNA was evaluated in 1% agarose gels stained with SYBR Green and ran at 100 V for 60 min.

### Genome Sequencing, Assembly, and Analysis

For the construction of sequencing libraries, (I) the genomic DNA was randomly fragmented by sonication; (II) DNA fragments were end polished, A-tailed, and ligated with the full-length adapters of Illumina sequencing, and followed by further PCR amplification with P5 and indexed P7 oligos; and (III) the PCR products as the final construction of the libraries were purified with AMPure XP system (Beckman Coulter Inc., Indianapolis, IN, USA). Sequencing library size distribution quality control was performed with an Agilent 2100 Bioanalyzer (Agilent Technologies, CA, USA) and quantified by real-time PCR (to meet the criteria of 3 nM). Whole genomes were sequenced using Illumina NovaSeq 600 using the PE 150 strategy at the America Novogene Bioinformatics Technology Co., Ltd.

Genomes were assembled using the Assembly HiSeq Pipeline, a SnakeMake pipeline to assemble sequencing data produced by Illumina (7). The pipeline integrates different quality control tools like FastQC (31) to analyze and visualize read quality, Adapter-Removal v2 (32) for removing sequencing adapters, and KmerStream (33) for computing k-mer distribution. For the genome graph construction, two main assemblers were used: Edena V3 (34) and Spades 3.9.1 (35); CD-HIT (36) and Unicycler (37) were used to optimize and integrate the assemblies previously produced. Whole-genome annotation was performed with RAST (38) and Prokka (39). To predict and reconstruct individual plasmid sequences in the genome assemblies, we used MOB-recon (40). Finally, QUAST (41) computed assembly quality metrics and each individual genome phylogenetic affiliation was confirmed through JSpeciesWS web tools (42) using the contigs generated by the assemblies. The whole genomes shotgun projects have been deposited to DDBJ/ENA/GenBank. Table 3 contains the main characteristics of the genomes.

The genomes were uploaded to the RAST (38) annotation server to identify the subsystem of each genome, obtaining information of genes related to different functions, including virulence, pathogenicity, plasmids and antibiotic resistance. Also, a pathogenicity and a virulence analysis were made for each genome through the Pathogen-Finder tool and the VirulenceFinder tool in the Center for Genomic Epidemiology. In addition, a plasmid identification was made through the PlasmidFinder tool also in the Center for Genomic Epidemiology. Finally, the identification of the resistomes through the CARD database Resistance Gene Identifier tool was made (43). There was a total of seven different species to describe.

Bioinformatic analyzes of the multi-resistant genomes started with Resistance Genes Identifier (RGI) with the CARD protein database (43) and ResFinder-4.0 (44) to predict the resistance genes. Plasmid detection was conducted through the MOB-suite (40) and PlasmidFinder-2.1 (45). For pathogenicity classification of each of the strains, PathogenFinder-1.1 (46) was utilized. VirulenceFinder-2.0 (47) was used to determine the virulence factors of each genome. The serotypes of the E. coli genomes were determined using SerotypeFinder-2.0 (48), and the number of mobile elements were determined by MobileElementFinder (49).

All genomes were properly deposited on NCBI’s SRA database with assessions: JAGJVC000000000,JAGJVD000000000,JAGJVE000000000,JAGJVF000000000, SAMN18612536, SAMN18612543, SAMN18612605.

## ACKNOWLEDGMENTS

This research project was successfully conducted thanks to the support provided by the Research Vice-Rectory and the Deanship of Basic and Environmental Sciences at Instituto Tecnologico de Santo Domingo.

The Federal University of Para team was support by the Coordenação de Aperfeiçoamento de Pessoal de Nível Superior - CAPES, Conselho Nacional de Desenvolvimento Científico e Tecnológico - CNPq, Pró-reitoria de Pesquisa e Pós-graduação(PROPESP)-UFPA and Pró-Reitora de Relações Internacionais (PROINTER)-UFPA

This research was partially funded by Fondo Nacional de Innovación y Desarrollo Científico y Tecnológico (FONDOCYT) of Miniterio de Eduacion Superior Ciencia y Tec-nología (MESCyT) grant number 2018-2019-2B4-157.

